# The Aging Slow Wave: A Shifting Amalgam of Distinct Slow Wave and Spindle Coupling Subtypes Define Slow Wave Sleep Across the Human Lifespan

**DOI:** 10.1101/2020.05.28.122168

**Authors:** Brice V. McConnell, Eugene Kronberg, Peter D. Teale, Grace M. Fishback, Rini I. Kaplan, Stefan H. Sillau, Angela J. Fought, A. Ranjitha Dhanasekaran, Brian D. Berman, Alberto R. Ramos, Rachel L. McClure, Brianne M. Bettcher

## Abstract

**Study Objectives:** Slow wave and spindle coupling supports memory consolidation, and loss of coupling is linked with cognitive decline and neurodegeneration. Coupling is proposed to be a possible biomarker of neurological disease, yet little is known about the different subtypes of coupling that normally occur throughout human development and aging. Here we identify distinct subtypes of spindles within slow wave upstates and describe their relationships with sleep stage across the human lifespan.

**Methods:** Coupling within a cross-sectional cohort of 582 subjects was quantified from stages N2 and N3 sleep across ages 6-88 years old. Results were analyzed across the study population via mixed model regression. Within a subset of subjects, we further utilized coupling to identify discrete subtypes of slow waves by their coupled spindles.

**Results:** Two different subtypes of spindles were identified during the upstates of (distinct) slow waves: an “early-fast” spindle, more common in stage N2 sleep, and a “late-fast” spindle, more common in stage N3. We further found stages N2 and N3 sleep are composed of two discrete subtypes of slow waves, each identified by their unique coupled-spindle timing and frequency. The relative contribution of coupling subtypes shifts across the human lifespan, and a deeper sleep phenotype prevails during old age.

**Conclusions:** Distinct subtypes of slow waves and coupled spindles form the composite of slow wave sleep. Our findings support a model of sleep-dependent synaptic regulation via discrete slow wave/spindle coupling subtypes and advance a conceptual framework for the development of coupling-based biomarkers in age-associated neurological disease.

**Statement of Significance:** Slow waves of nonrapid eye movement sleep couple with sleep spindles in a process hypothesized to support memory functions. This coupling has recently gained interest as a possible biomarker of cognitive aging and onset of Alzheimer’s disease. Most studies have been limited by an assumption that all slow waves (and coupled spindles) are fundamentally the same physiological events. Here we demonstrate that distinct subtypes of slow waves and their coupled spindles can be identified in human sleep. A mixture of different slow wave and spindle subtypes shifts in composition during lighter versus deeper sleep, and aging favors the deep sleep subtypes. These data should inform any future attempts to use slow wave sleep as a biomarker or clinical interventional target.

## Background

Slow waves occur during stages N2 and N3 Non-Rapid Eye Movement (NREM) sleep and are associated with large-scale synchronization of neuronal populations.^1-3^ Spindles are generated and propagated via cortical-thalamic loops, with timing input from the reticular nucleus of the thalamus.^4,5^ In the process of aging, loss of slow waves and sleep spindles is highly correlated with cognitive decline, and abnormal slow wave neuronal circuitry is implicated in the pathogenesis of Alzheimer’s disease^6^.

The dynamic coordination of slow waves and spindles provides a fundamental framework for synaptic regulation during sleep-dependent memory consolidation.^7,8^ During slow wave sleep, reactivation of memory circuits is coordinated by spindle oscillations.^9-11^ Higher frequency oscillations, termed ripples, are nested within spindles and accompany specific patterns of hippocampal memory reactivation.^12-15^ Together, slow waves, spindles and ripples maintain a triple coupling relationship that supports synaptic plasticity and remodeling of cortical networks during sleep.^16-19^

Timing aspects of slow wave and spindle coupling are compromised among aging adults, and abnormal coupling correlates with brain atrophy and deficits in sleep dependent memory performance.^8,20,21^ Loss of coupling integrity has further been demonstrated to correlate with tau deposition among cognitively intact aging adults, raising the prospects of slow wave and spindle coupling as a possible functional biomarker of early Alzheimer’s pathogenesis.^22^ Given the role of spindles in regulation of synaptic function,^8,23,24^ an understanding of the pathophysiology of spindle coupling abnormalities may also provide fundamental mechanistic insights into the proposed role of sleep in protecting against neurodegenerative disease.^6,25,26^

Understanding how the timing aspects of coupling might subserve or modify pathological outcomes is currently limited by an incomplete characterization of the key coupled-event subtypes that occur during neurodevelopment and the aging process. Original descriptions of coupled spindles identified slow and early-fast subtypes, occurring before and after the trough of the slow wave downstate, respectively.^27,28^ More recently, human intracranial recordings have also identified additional coupling events pre-downstate, trough termed theta bursts, that are proposed to coordinate hippocampal ripples^29,30^ and may trigger slow wave events^31^. During the slow wave upstate, “faster” and “slower” spindle subtypes have also been observed within intracranial recordings, with the faster subtype occurring before slower spindles in temporal relation.^31,32^ Little is known about what the timing aspects of these events may signify in the greater context of human sleep and cognitive functions.

Slow wave subtypes have also been proposed, and a model of global slow waves versus local slow waves postulates different neuroanatomical generation of these slow wave subtypes.^33,34^ Global slow waves are hypothesized to be generated via a frontal lobe process, with a predominance in stage N2 sleep. These waves are typically more interspersed rather than tightly clustered and are proposed to include the classically described K-complexes.^33,34^ In contrast, local slow waves are thought to be generated more widely throughout the brain and occur more commonly during stage 3 sleep.^33,34^ Recent evidence from murine sleep further advances a conceptual model in which a global subtype of slow wave/spindle coupling favors synaptic potentiation, while a distinct local slow wave/spindle coupling subtype drives synaptic downscaling.^35^ Despite accumulating evidence that multiple subtypes of slow waves occur in human sleep, it remains unclear how different subtypes of spindles might couple to these different slow waves.

Based on the conceptual framework that coupling integrity is integral to human memory function and a potential biomarker of neurodegenerative disease, we sought to expand the fundamental understanding of coupling physiology by characterizing distinct subtypes of slow wave and spindle coupling across the human lifespan. We tested the hypothesis that subtypes of spindles preferentially occur during different sleep stages, and that predominant coupling subtypes shift throughout neurodevelopment and aging. Further, as a critical step toward elucidating the underlying physiological basis of spindle coupling subtypes, we examined whether coupled spindles could be utilized as an “identification tag” to sort out a mix of different slow wave event subtypes from NREM sleep.

Drawing from 582 subjects aged 6-88 years old from the Cleveland Family Study,^36-39^ our analyses demonstrate that two subtypes of coupling, “early-fast” and “late-fast,” can be identified during the upstates of slow waves. The early-fast coupling subtype preferentially occurs in stage N2 sleep during early life, with a gradual transition to a higher proportion of late-fast coupling subtype as age advances. Meanwhile, the late-fast coupling subtype is the predominant coupling event in stage N3 sleep after early childhood through the later stages of life. We further provide key mechanistic insights by demonstrating that the coupling of spindles can be used to identify discrete populations of slow waves that differ in their association with stage of sleep. Taken together, our findings advance the fundamental conceptual framework of slow wave and spindle coupling and provide key methodological advancements for the development of coupling as a functional biomarker in human neurological disease.

## Methods

### Participants

Participant data was accessed from the Cleveland Family Study (CFS) via the National Sleep Research Repository.^36-39^ The CFS was conducted as a longitudinal study of sleep apnea and included 2284 individuals from 384 families over 16 years of follow up. All data were collected with written informed consent under protocols approved by a local institutional review board at each institution. Overnight polysomnography was utilized from visit 5 study data for 735 subjects (Figure 1). Data quality was rated by sleep staging experts per standardized study protocol, and EEG recordings with less than 95% overall quality score were excluded from our analysis (n=132). Recordings with problems noted in differentiating N2 and N3 sleep were also excluded (N=4), as well as a diagnosis of Parkinson’s disease (n=1), and participants having taken medications for sleep within three days up to the overnight polysomnography (n=9). An additional seven subjects we excluded due to data quality problems during processing, resulting in a total of 582 subjects for analysis.

**Figure 1:**
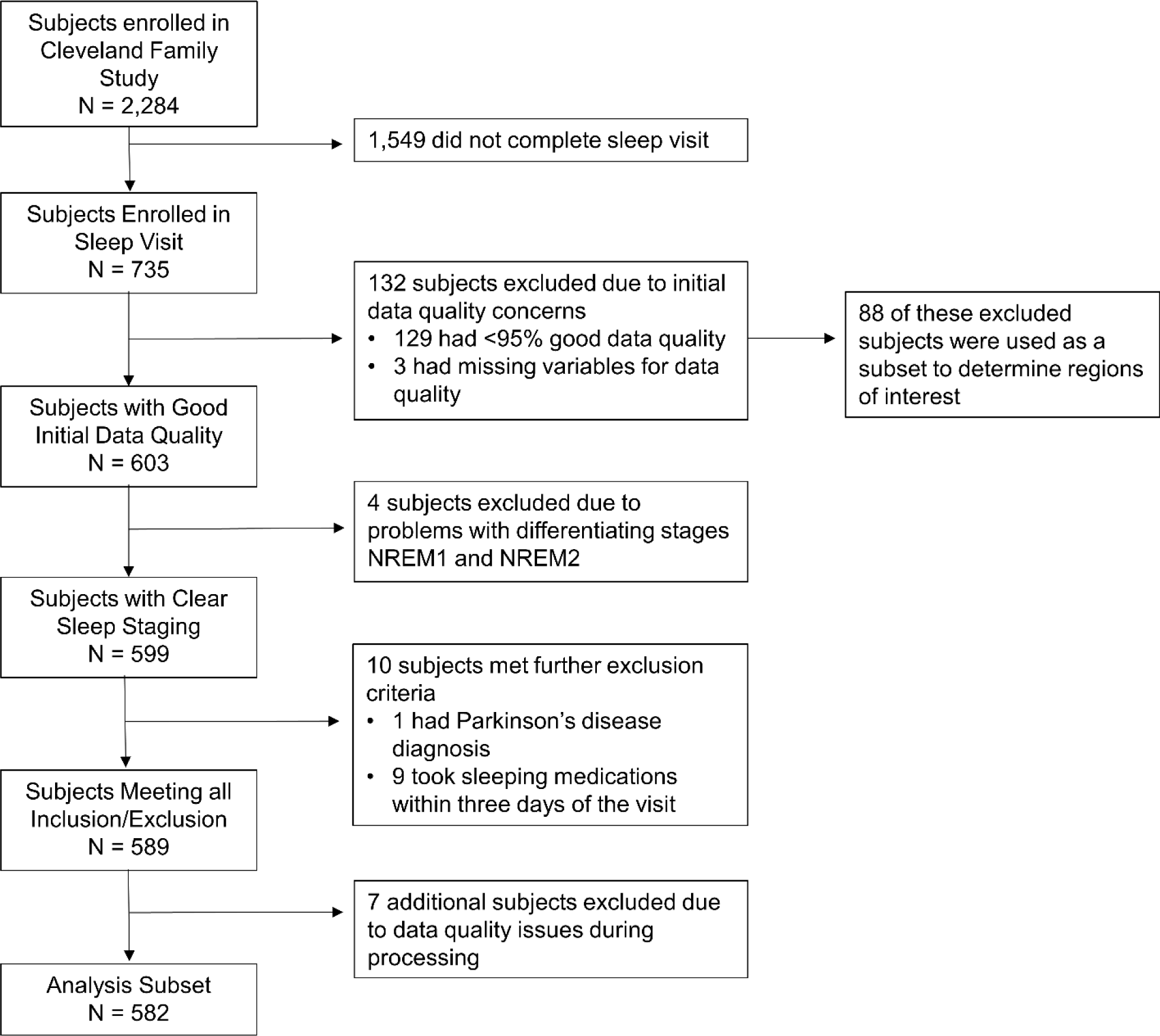
Subject selection flow chart diagram details exclusion criteria

### EEG Acquisition

Laboratory-based polysomnography was performed using a 14-channel Compumedics E-Series System (Abbotsford, Australia). EEG was obtained using gold cup electrodes at a sampling rate of 128 Hz (filtered high pass 0.16 Hz and low pass 64 Hz). Scalp electrodes C3 and C4 were referenced to positions A2 and A1, respectively. Recordings were scored by trained technicians at a centralized reading center at Brigham and Women’s Hospital in accordance with uniform study protocols and AASM criteria.^40^

### Spectral Analysis

All EEG processing was performed in MATLAB (R2018b; the Mathworks, Inc., Natick, MA) using additional toolboxes including EEGlab^41^ and FieldTrip.^42^ Multitaper spectral analysis was performed on raw EEG data using discrete prolate slepian sequences as described by Prerau et al.^43^ Briefly, data were epoched to 30 second segments with 85% overlap. A frequency range of 0.5 to 30 Hz with a resolution of 0.01 Hz were used to obtain spectral estimates. In total, 29 tapers were created to achieve a frequency smoothing of ±0.5 Hz.

### Slow Wave Detection

Data for slow wave detections was processed from electrodes C3 and C4 from stages N2 and N3 of slow wave sleep by AASM criteria.^40^ Slow waves from stage N2 were identified for all subjects. Due to lack of stage N3-specific slow wave events in subjects with minimal or no N3 sleep, we set a minimum of 5% time in N3 (from total sleep) for detection of N3-specific slow waves. Epochs with un-scorable data were excluded from analysis.

Preprocessing for high throughput slow wave identification at the population level was performed via automated artifact reduction (n=661 subjects, including those used exclusively for ROI determination). Automated management of high amplitude artifacts was accomplished via exclusion of regions of range greater than 900 µV. Remaining raw EEG segments of at least three seconds were detrended; these sections were cut for detrending when the final three seconds exceeded a range of 900 µV. A small cohort was processed for further slow wave and spindle characterization with additional manual visual artifact removal (n=10 subjects). Manual artifact removal was performed within EEGLAB in 5 second bins with visual inspection and removal of artifacts.

Parameters for slow wave detection were based on prior slow wave/spindle coupling reports.^20,44^ Sequentially, EEG data was detrended to remove any remaining start-end mismatch that would cause ringing and then band-pass filtered in a forward and backward direction using a 6^th^-order Butterworth filter between 0.16-4 Hz. Zero crossings were identified to detect negative and positive half-waves, and slow wave events were identified when the half-wave pair zero crossings occurred between 0.5 to 1 sec. Minimal and maximal half-wave amplitudes were measured from the zero crossing, and slow waves with both positive and negative maximum amplitudes in the top 40% of all waves were selected for subsequent coupling analysis. We rejected all zero crossing pairs with peak/trough amplitudes exceeding four standard deviations from the mean min/max zero crossing pair amplitudes for each subject. An additional visual inspection was performed on each individual recording’s slow wave average composite to ensure correct channel polarity.

### Time-Frequency Analysis

Time-frequency wavelet plots of EEG data were produced to measure coupling of spindles by adapting established methods.^20,27^ A Morlet-wavelet transformation (8 cycles) was applied to unfiltered EEG regions of 5-seconds for slow wave and baseline segments between 4 Hz and 20 Hz in steps of 0.25 Hz. For each 5-second region surrounding the centered trough of a slow wave, a baseline region of matching time length was selected from the interval immediately preceding a run of slow wave. The mean of these baseline regions was used to normalize the resulting amplitude of the mean Morlet-wavelet transformation of all slow-wave event regions. A manual inspection of each time-frequency map was performed to remove any time-frequency maps from analysis that contained ringing artifacts secondary to bandpass filtering (<7% excluded total; <2.5% excluded from any 10 year age increment).

### Region of Interest (ROI) Selection

Regions within the time-frequency wavelet plots were selected for quantitative analysis utilizing a subset of 88 subjects (ages 15-64) from the Cleveland Family Study (Supplementary Table 1). These subjects were noted to have between 75-95% good EEG data quality, and thus were excluded from our main analysis cohort and restricted to our efforts to define the boundaries of ROIs. An average composite of channel C3 was created for all 88 subjects, and four regions of interest were manually selected for subsequent statistical analysis of the main cohort. Theta bursts were defined by a ROI defined between 4 Hz to 5.5 Hz at −0.7 sec to 0.5 sec from the average slow wave trough, plus an additional region 5.5 Hz to 7 Hz at −0.5 sec to 0.4 sec from the trough. Pre-trough slow spindle ROI was defined as 7 Hz to 11 Hz, and −0.4 sec to 0.2 sec from the average trough. Early-fast spindle ROI was defined as 14 Hz to 17 Hz and 0.1 sec to 0.7 sec from the average trough. Late-fast spindle ROI was defined as 10 Hz to 14 Hz, and 0.4 sec to 1.1 sec from the average slow wave trough (Supplementary Figure 1).

### Slow Wave Separation by Region of Interest Spindle EEG Power

Normalized EEG power from regions of interest corresponding to the early-fast and late-fast spindles were utilized to sort individual slow waves. Each slow wave was sorted by calculating a ratio of the EEG power values in the corresponding early-fast and late-fast spindle ROIs. Given unknown prevalence of each slow wave subtype, here we modeled an approximate 50/50 mix of slow wave subtypes based on the early-fast spindle percentage (early-fast/[early-fast+late-fast]) values that were estimated to range from 51.2% to 48.6% across the study population. Using these study-specific parameters, all of the identified slow waves were sorted by top 50% of early-fast spindle ROI power ratio (early-fast/late-fast) vs top 50% of late-fast spindle power (late-fast/early-fast).

### Slow Wave Separation by Co-Identification of Spindle Events and Timing Alignment

Parameters for spindle detection were based on prior slow wave/spindle coupling reports27,45 Sequentially, artifact free EEG data was detrended and bandpass filtered in a forward and backward direction using a 3rd-order Butterworth filter between 10-14 Hz for late-fast spindles, and between 14.5-17.5 Hz for early-fast spindles. Next the upper spindle envelopes were calculated and an amplitude threshold of 75% percentile of the root mean squared value, with a length window of 0.5 sec to 3.0 sec was used to define spindle events. An absolute threshold of 40 microvolts in range was used to eliminate artifacts and then only waves within 8 standard deviations from the mean amplitude values were used in further analysis. Each spindle event was indexed with respect to adjacent spindle events to determine their ordering. Slow wave events coupled to these populations of spindles were identified by their co-localization within 0.5 to 1.5 sec from the trough of a slow wave. These individual slow waves that coupled to each spindle type (late-fast and early-fast) in the time domain were then selected for additional analysis as coupled events of interest.

### Statistical Analysis

All statistical modeling was performed with SAS v9.4 (SAS Institute Inc., Cary, NC, USA). Normalized EEG power values were obtained from time-frequency wavelet plots within each of the four ROIs (early-fast, late-fast, slow, and theta) that were defined *a priori*. The ROI values for each measurement were statistically weighted by the number of slow wave events that were utilized to calculate each corresponding value. To evaluate the relationship with power from each ROI and age, splines were used to find knots at ages 20 and 60 for characterizing different neurologic stages of life (childhood/adolescence ages 6-20, young adulthood through late-adulthood ages 21-60, and late age 61 and above). Then two main power outcomes were created, 1) early-fast divided by the sum of early-fast and late-fast and 2) late-fast divided by the sum of all 4 ROIs (this second analysis was only used to calculate relationships with age as different sleep stage baseline EEG segments confound comparisons between sleep stages due to frequency leakage in the theta range during stage N3 sleep).

Mixed models were used to account for repeated measures from combinations of the electrode (C3 and C4) and stage (N2 and N3), adjusting for sex, sleep apnea (using a binary threshold at 15 apneas and hypopneas with >= 3% oxygen desaturation or arousal per hour of sleep), and age as piece-wise continuous with flex points determined by the knots at 20 and 60, driven by the spline analysis. Interactions with stage, electrode, and electrode and stage were included for each covariate (early-fast, late-fast, slow, and theta). In each model, we accounted for familial clustering by including family identification number as a random effect. An additional model, stratified by electrode, similarly accounted for repeated measures with the same covariates and interactions, and in this case outcomes were log-transformed normalized EEG power with a covariate specifying the corresponding ROI (early-fast, late-fast, slow, and theta). Linear combinations of the model parameters were constructed and tested with T and F tests, with estimates and confidence intervals for individual linear combinations. Notably, statistical modeling was performed for both C3 and C4 electrodes, and here we report C3 because we did not identify statistically significant differences when estimates were compared between electrodes for the main outcomes.

## Results

### Stage N2 and Stage N3 Slow Waves Demonstrate Distinct Compositions of Slow Wave and Sleep Spindle Coupling

Given that slow wave activity is described to have different functions between stage N2 and stage N3 sleep,^45^ we sought to characterize populations of slow waves from each sleep stage independently using surface EEG overnight recordings from subjects within the Cleveland Family Study.^36-39^ Notably, the distinction between sleep stages is imperfect, and most often a mixing of lighter and deeper sleep occurs, particularly in transitions between sleep stages.^46,47^ Further, individual variability of slow waves and spindles has been demonstrated with regard to frequency and timing aspects of coupling.^48^ Nonetheless, dividing sleep into stages N2 and N3 sleep demonstrated that distinct slow wave and spindle coupling features can be distinguished among the composition of slow waves from stages N2 and N3 sleep (Illustrated in Figure 2).

**Figure 2:**
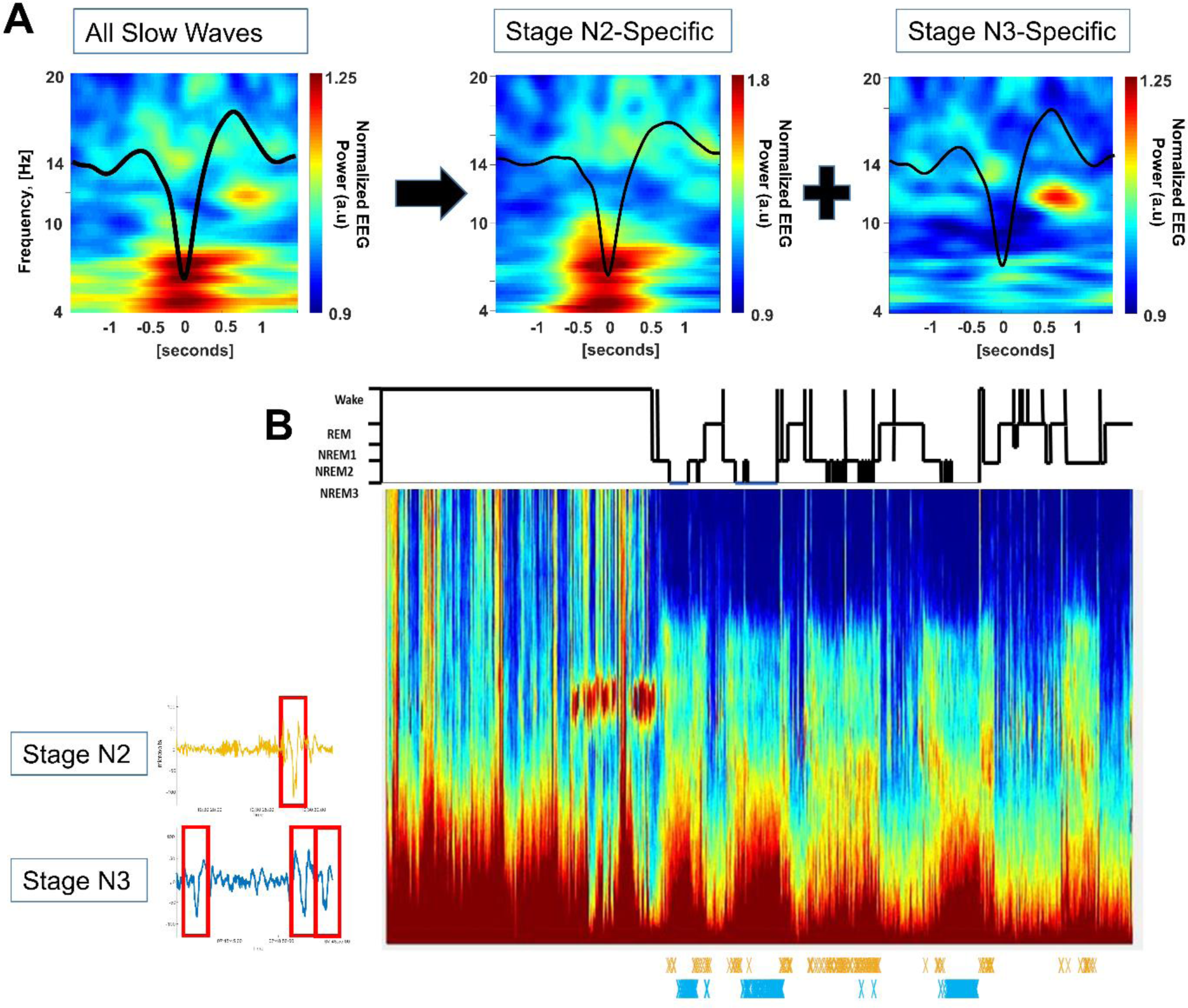
Stage-specific slow wave detection example. (A) Time-frequency wavelet plots from a single subject (age 20) demonstrates distinct coupling between N2 and N3. (B) An illustrative example of slow wave detections from a single subject. Individual slow wave detections are marked with “X” and illustrated in temporal relationship with a full night multitaper spectrograph for sleep stages N2 and N3 separately (orange = N2-specific detections; blue = N3-specific detections).

To better understand the composition of these coupling events across the lifespan, we first established *a priori* regions of interest (ROIs) using overnight EEG recordings from a cohort of 88 subjects, ages 15-64 (these subjects were noted to have 75-95% good EEG data quality, and thus were only used to define boundaries of ROIs for our statistical model; Supplementary Table 1; Supplementary Figure 1). Our analyses focused on four regions of interest as peri-slow wave coupling events (Figure 3A). These regions include pre-slow wave trough activity corresponding to possible theta burst (4-7 Hz)^29-31^ and slow spindle activity (7-11 Hz).^27,49,50^ The post slow wave trough demonstrates coupling activity consistent with previously described fast and slow spindles of the slow wave upstate,^31,32^ herein referred to as early-fast spindles (14-17 Hz) and late-fast spindles (10-14 Hz). We then applied these slow wave characterization techniques and ROI boundaries to a cohort of 582 subjects (ages 6-88; Table 1) to examine population-level differences in slow wave and spindle coupling.

**Table 1.** Demographics of the study population.

| *Characteristic* | *N* | *N (%) or Median (IQR)* |
| --- | --- | --- |
| Age | 582 | 43.01 (22.20-54.79) |
| Sex | 582 |  |
| - Female |  | 321 (55.15) |
| - Male |  | 261 (44.85) |
| Race | 582 |  |
| - White |  | 237 (40.72) |
| - Black |  | 324 (55.67) |
| - Other |  | 21 (3.61) |
| Apnea Hypopnea Index | 582 | 4.71 (1.57-14.14) |
| Apnea Hypopnea | 582 |  |
| - Less than 15 |  | 446 (76.63) |
| - 15 or greater |  | 136 (23.37) |
| % Time in Stage 1 | 575 | 4.13 (2.57-6.03) |
| % Time in Stage 2 | 582 | 55.30 (47.70-64.20) |
| % Time in Stage 3 | 582 | 19.85 (12.10-28.20) |
| % Time in REM | 580 | 18.25 (14.00-22.60) |

**Figure 3:**
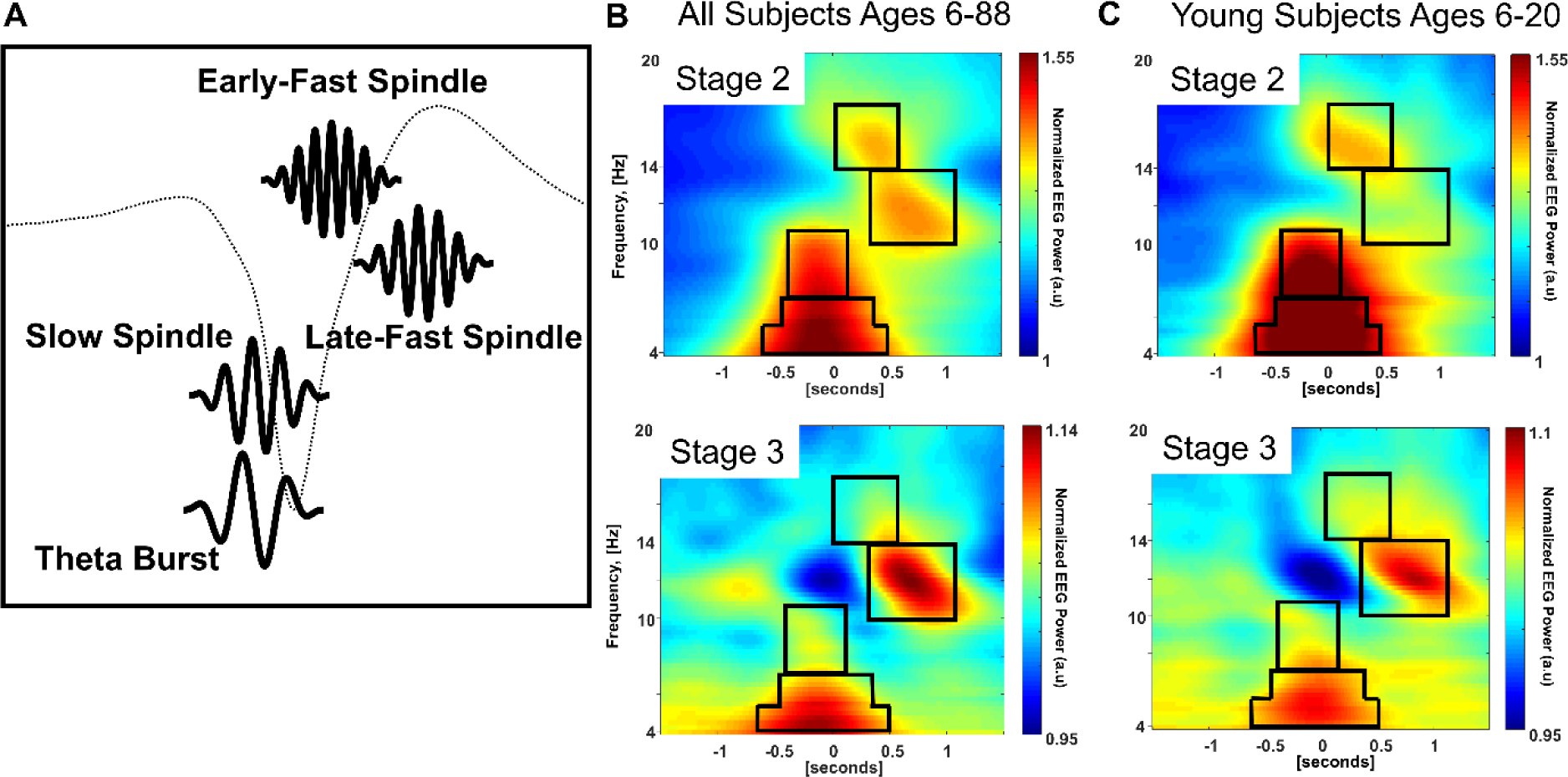
Region of interest selection. (A) Four regions of interest were identified a priori (using a separate cohort) as peri-slow wave coupled events. (B) The average of wavelets across all age groups demonstrates differences in relative intensities between stages N2 and N3 wavelets. (C) Average wavelets of subjects 20 years old or less demonstrates a relatively higher proportion of early-fast spindles compared to late-fast spindles in stage N2 sleep

Differences in relative intensities of these ROIs are apparent between stages N2 and N3 sleep (Figure 3B). To quantify differences between sleep stages, we utilized statistical modeling to calculate adjusted values for each of the regions of interest that define these peri-slow wave events. We utilized relative calculations between ROIs to make meaningful comparisons between sleep stages and subject age despite differences in ROI size and non-linear changes in EEG power across frequencies. Our primary focus was a calculation of early-fast spindle power relative to total post-trough spindle coupling by percentage (early-fast spindle/[early-fast spindle + late-fast spindle]). Given the findings by Muehlroth et al. demonstrating relationships between post-trough spindle frequency and timing changes with sleep-dependent memory consolidation,^20^ we hypothesized that these post-trough spindle ROIs could be used to differentiate features of slow wave populations obtained from stages N2 and N3 sleep.

### The Greatest Differences between Stage N2 and Stage N3 Coupling Are Among the Youth

We observed that the differentiation of sleep stage-specific slow wave and spindle coupling features is dependent on age, and the most appreciable distinction between stages N2 and N3 was observed among young adults and children. When averaged across all ages, stage N2 sleep contains both early-fast and late-fast spindle coupling post-slow wave trough (Figure 3B). Younger subjects, here illustrated by averaging subjects between 6-20 years old, demonstrate a distinction between a higher percentage of early-fast spindles in N2 versus N3 sleep (Figure 3C; p <0.001).

### Early-Fast Spindle Coupling is Prevalent in the Younger Years of Life

Tracking with younger life brain maturation from adolescence to young adulthood, early-fast spindle coupling comprises a relatively high percentage of post-slow wave coupling during stage N2 sleep (Figure 4A). To illustrate the prevalence of early-fast spindle coupling in early life (i.e., defined as ages 6-20), we utilized mixed effects models to estimate mean stage N2 early-fast spindle ROI percentages at age 8 years old (51.16%, 95% CI 50.23% to 52.09%), age 13 years old (50.83%, 95% CI 50.27% to 51.39%), and 18 years old (50.51% 95% CI 50.11% to 50.90%). Subjects within this neurodevelopmental stage also demonstrate a relatively high percentage of early-fast spindle activity in stage N3 sleep (Figure 4B). Stage N3 early-fast spindle ROI percentages from the statistical model were estimated at age 8 years old (50.09%, 95% CI 49.69% to 50.50%), age 13 years old (49.84%%, 95% CI 49.57% to 50.11%), and 18 years old (49.58% 95% CI 49.34% to 49.83%).

**Figure 4:**
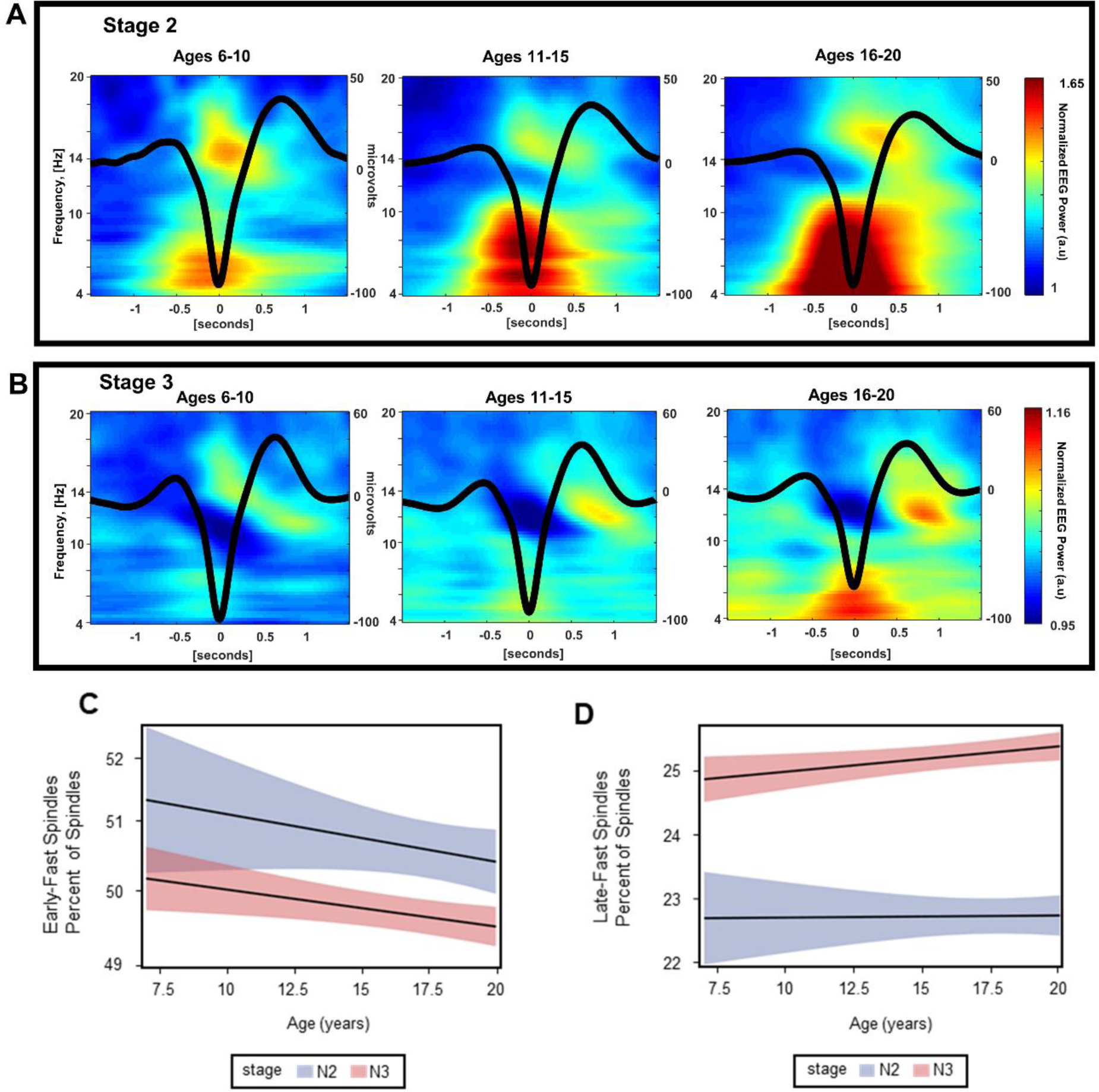
Changes in spindle coupling during childhood and adolescence. (A) Sleep stage N2-specific time-frequency wavelet plots visually illustrate higher percentage of early-fast spindles. (B) Sleep stage N3-specific time-frequency wavelet plots visually illustrate a transition from higher-to-lower percentage of early-fast spindles. (C) Statistical modeling with 95% confidence interval estimation for the percentage of early-fast spindle ROI through childhood to young adulthood (ages 6-20) (D) Statistical modeling with 95% confidence interval estimation for the percentage of late-fast spindle ROI through childhood to young adulthood (ages 6-20)

### Young Adulthood through Late Life Demonstrates a Shift in N2 Coupling with Relative Preservation of N3 Coupling

We next expanded the scope of age-related coupling changes across the adult portion of the human life cycle (i.e., ages 21 and above). Aging through midlife to late life is associated with a progressive loss of pre-trough theta burst and slow spindle activity in N2 sleep (Illustrated from ages 21-80 in Figure 5A; Statistical model in Supplementary Figure 2). The post-slow wave percentage of early-fast spindles of stage N2 sleep also progressively declines with age in midlife. Focusing on young adulthood through late adulthood, estimates from our mixed model project a rate of decline at −0.28% early-fast spindle ROI percentage per 10 years from ages 21-60 years old (p <0.001, 95% CI −0.43% to −0.13%). The shift to a higher percentage of late-fast, post-slow wave spindles in stage N2 sleep occurs early in young adulthood, and these late-fast spindles become increasingly predominant post-trough with advancing age (Figure 5A). This relative gain in late-fast spindle ROI power was estimated from our statistical model as a percentage late-fast spindle out of the total ROIs values (late-fast spindle/[early-fast spindle + late-fast spindle + theta burst + slow spindle]). The rate of late-fast spindle gain was 0.66% per 10 years from ages 21-60 years old (p <0.001, 95% CI 0.55% to 0.77%).

**Figure 5:**
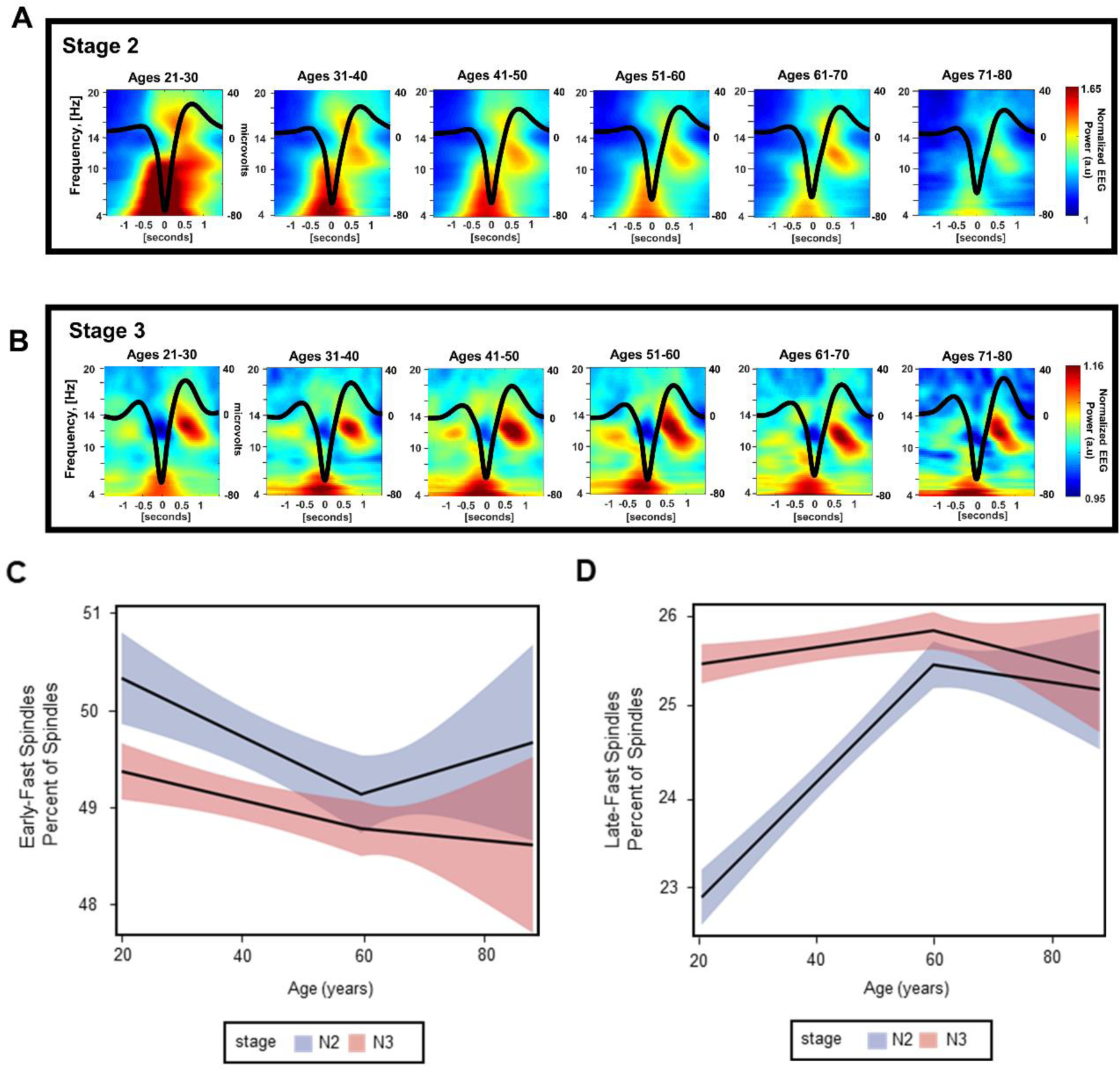
Changes in spindle coupling during young adulthood through late age. (A) Sleep stage N2-specific time-frequency wavelet plots visually illustrate higher percentage of early-fast spindles. (B) Sleep stage N3-specific time-frequency wavelet plots visually illustrate a transition from higher-to-lower percentage of early-fast spindles. (C) Statistical modeling with 95% confidence interval estimation for the percentage of early-fast spindle ROI through childhood to young adulthood (ages 21-88) (D) Statistical modeling with 95% confidence interval estimation for the percentage of late-fast spindle ROI through childhood to young adulthood (ages 21-88)

Stage N3 sleep is notable for relatively stable late-fast spindle power (Figure 5D). Between ages 21-60 years old, the percentage of late-fast spindle change was estimated at 0.09% increase per 10 years of age (p=0.015, 95% CI 0.02% to 0.17%). Early-fast spindles demonstrate a moderate decline during stage N3 sleep with age (Figure 5C), with an estimated −0.14% change in early-fast spindle percentage per 10 years from ages 21-60 years old (p=0.004, 95% CI - 0.23% to −0.04%). Estimated rates of change within early-fast and late-fast spindle percentages during stages N2 and N3 sleep above age 60 were not statistically significant.

### Slow Wave Subtypes Can Be Separated by Their Coupling to Early-Fast and Late-Fast Spindles

We next employed a mechanistic approach to better understand the physiological basis for differences in early-fast and late-fast spindle coupling that we observed at the cohort level. We tested the hypothesis that not all slow waves from our NREM sleep data are fundamentally the same physiological process (i.e. there are distinct subtypes of slow waves that couple with different spindles). We specifically tested whether early-fast and late-fast spindles can be used as an “identification tag” to find and sort distinct subtypes of slow waves.

We focused this approach on a subset of subjects aged 21 years old (n=10), selected *a priori*, to examine an adult sample while minimizing potential confounds due to the age-related shifts in spindle coupling observed across the study population. Starting with a set of heterogeneous slow wave/spindle couples from combined N2 and N3 sleep, we used our defined ROIs to threshold normalized EEG power and separate apart discrete subtypes of slow waves with relatively high coupling activity in each respective ROI (Figure 6A,B,C).

**Figure 6:**
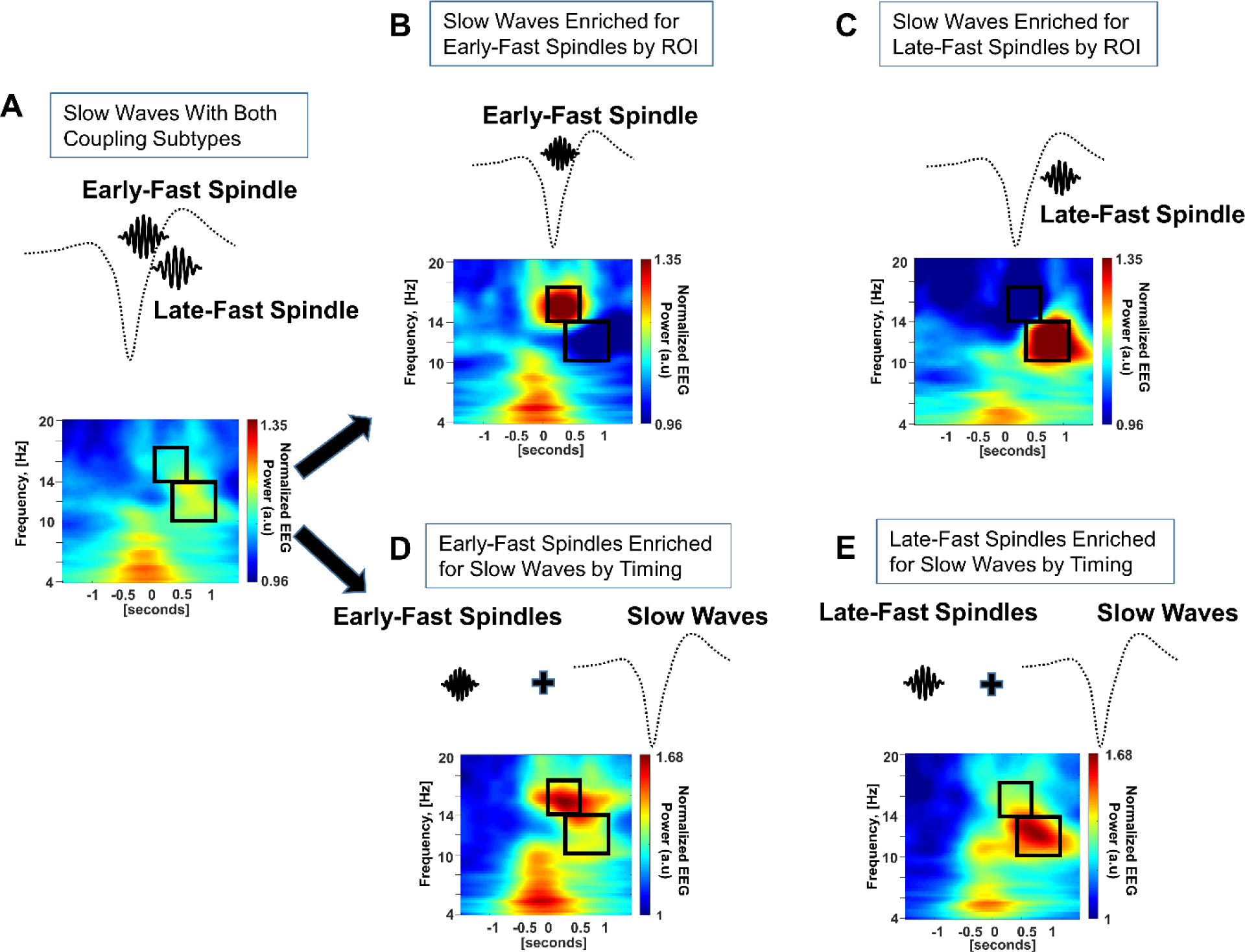
Two methods are used to cross-validate separation of distinct slow wave and spindle coupling pairs (Data averaged from 10 subjects). The original composite of subtypes includes both subtypes of slow wave and spindle coupling (A). Slow waves coupled to early-fast and late-fast spindles were enriched by their relatively high power within the respective ROIs (B and C). Early-fast and late-fast spindle events were first identified independently, then used to identify their respective coupled slow wave events

In order to cross-validate this sorting technique and minimize circularity, we next sorted slow wave and spindle coupled pairs using a different signal processing method that does not depend on the ROI values (i.e. re-creating the spindle “identification tags” without the dependence on time-frequency wavelet data ROIs). We again utilized the EEG data from combined N2 and N3 sleep from the same ten subjects aged 21 years old. The early-fast and late-fast spindles from each recording were first identified as unique events from the raw EEG data. Notably, the frequencies of early-fast and late-fast spindles overlap around 14 Hz, therefore we moved the bandpass window to 14.5 - 17.5 Hz for early-fast spindles to reduce misidentification that might occur due to application of bandpass filtering EEG near this shared frequency range (late-fast spindles were still identified between 10 - 14 Hz). Next, early-fast and late-fast spindle events were assessed for co-occurrences within a time window of −0.5 sec to 1.5 sec near a downstate trough of a slow wave event. Using this method to “re-couple” the early-fast and late-fast spindles with their respective slow waves once again identified distinct populations of slow waves. Comparing the location of these spindles in time and frequency to the previously identified ROIs, this method produced striking agreement to our original approach with regard to both timing and frequency attributes (Figure 6D and E).

### Separated Slow Wave Subtypes Demonstrate Greater Late-fast Spindle Coupling During Stage N3 Sleep

Having observed that mixed populations of slow waves demonstrate greater late-fast spindle coupling during stage N3 sleep, we next sought to verify this finding among the isolated slow wave subtypes. Using the same 10 subjects aged 21 years old, we repeated our separation methods to sort each slow wave by their relative early-fast and late-fast spindle EEG power for stages N2 and N3 sleep independently. We again observed that stages N2 and N3 sleep contain a mix of both coupled slow wave subtypes (Figure 7A-F). Notably, the normalized EEG power within the late-fast spindle ROI is disproportionately high when compared to early-fast spindle ROI among the slow waves from stage N3 sleep (Figure 6 D and F).

**Figure 7:**
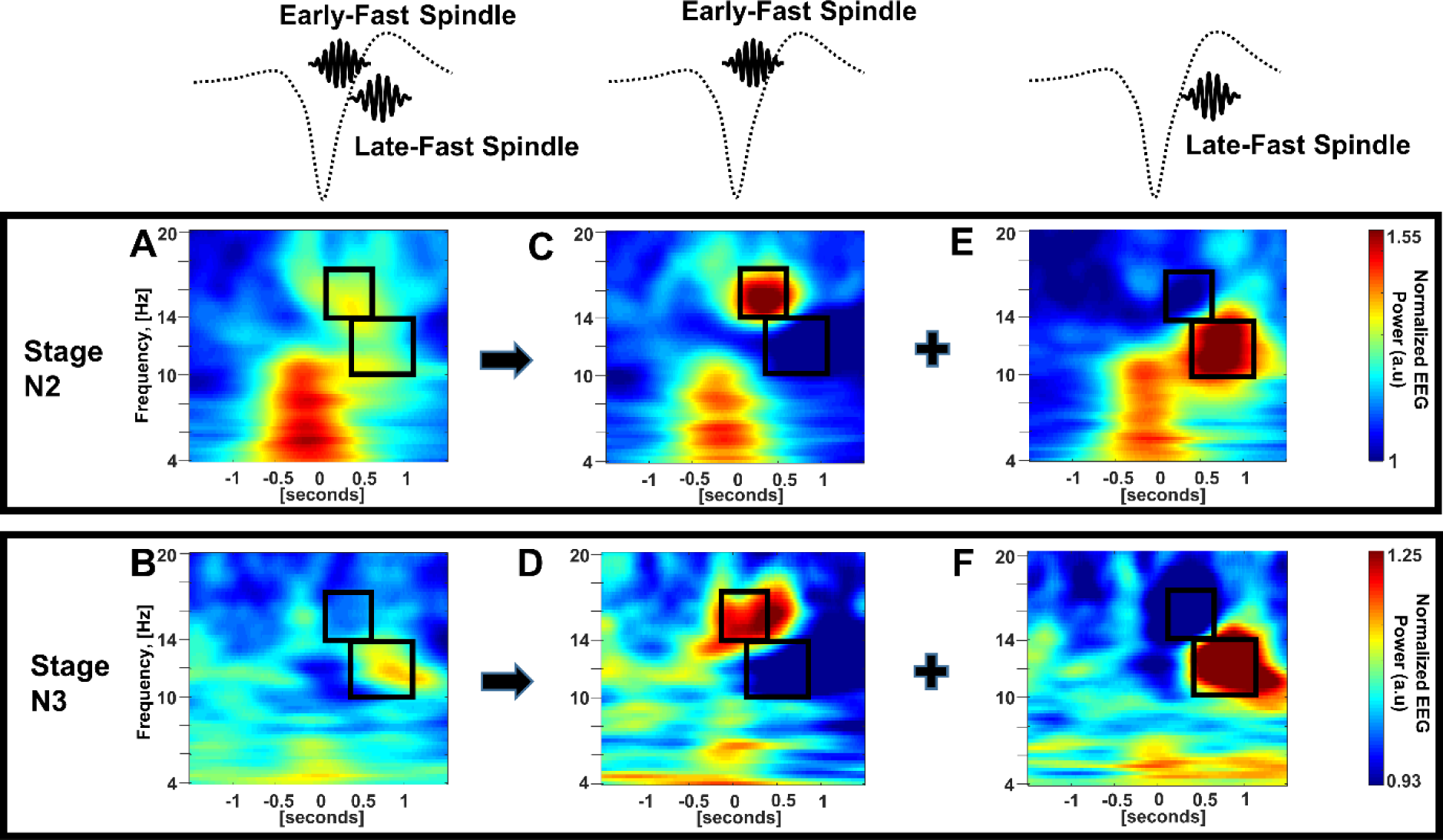
Sleep stage specific slow wave sorting (Data averaged from 10 subjects). Slow waves were identified by their coupling to early-fast or late-fast spindle subtypes by ROI and separated into two distinct slow wave populations from combined stages N2 and N3 sleep (A and B), stage N2 sleep (C and D), and stage N3 sleep (E and F).

## Discussion

Our results support a model in which distinct subtypes of spindle coupling extend beyond the “slow” and “fast” subtypes that were originally described.^27,28^ The existence of two distinct, temporally dissociated, spindles post-slow wave trough has been reported from intracranial recordings,^31,32^ and here we establish that indeed “early-fast” and “late-fast” spindle subtypes can be identified via surface EEG. The early-fast subtype occurs closer to the slow wave trough, at the beginning of the upstate, while the late-fast subtype is maximal closer to the peak of the upstate. We further report that both stage N2 and N3 sleep contain a composite of early-fast and late-fast coupling subtypes. Despite the presence of mixed subtypes, the relative amount of different coupling subtypes within each sleep stage can be reliably measured, is age dependent, and demonstrates a continuum across the human life span. Childhood and adolescence are associated with relatively higher early-fast spindle coupling, whereas the late-fast post-slow wave trough coupling subtype becomes increasingly predominant through mid-life and into the later stages of life. We further provide compelling mechanistic data demonstrating that the coupling of different spindle subtypes can be used to identify distinct slow wave subtypes during both N2 and N3 sleep.

The coordinated coupling of slow waves and sleep spindles provides a conceptual framework for understanding sleep’s role in synaptic regulation and memory consolidation.^18^ Among the most challenging gaps in this framework are the conflicting findings as to whether coupled spindles are responsible for potentiating vs downscaling synaptic strength.^51^ More current models that seek to reconcile these seemingly opposing findings propose that synapses are selectively downscaled or potentiated by distinct mechanisms during NREM sleep.^52,53^ Indeed, a study by Kim et al. recently demonstrated in murine sleep that two subclasses of slow wave and spindle coupling demonstrate opposing effects in synaptic potentiation vs downscaling. This study further reported that globally-occurring slow wave and spindle coupling is associated with synaptic potentiation, while the local subtype of slow wave and spindle coupling drives downscaling.^35^

While a human equivalent of this dual system of opposing synaptic regulation has not been identified, human NREM sleep does contain a mix of global and local slow wave events.^33,34^ Global events, also termed Type I slow waves, have been suggested to represent K complexes and are generated independent of local sleep pressure^33,34^. Local, Type II, slow wave events demonstrate a dynamic correlation to ongoing homeostatic sleep pressure and are thought to be “delta waves” of deeper stage N3 sleep^33^. Although our analyses are not able to attribute global versus local characteristics to slow wave and spindle coupling subtypes, we do observe a distinction between the prevalence of discrete coupling subtypes throughout lighter N2 and deeper N3 sleep. Future studies will be required to determine whether early-fast and late-fast subtypes of spindle coupling may have specificity to global or local slow wave event subtypes.

Changes in slow wave and spindle coupling relationships have been shown to correlate with performance on overnight memory consolidation, and age-dependent loss of coupling is associated with poorer performance on these memory tasks.^20,21^. Our analyses suggest that changes in this neuronal circuitry occur across the human lifespan. Childhood is marked by relatively higher early-fast spindle EEG power in both stages N2 and N3 sleep, with a progressive decline through adolescence, possibly reflecting neurodevelopmental changes in synaptic regulation. Further, we observe shifts in coupling as a progression from young adulthood through old age, and these changes may reflect a continuum sleep physiology’s contributions to well-documented lifelong changes in memory functions.^54^

Degradation of coupled spindle timing may signal neuronal dysfunction early in the pathogenesis of Alzheimer’s disease,^22^ raising prospects of developing a coupling-based functional biomarker of disease pathology. Our finding that distinct subclasses of slow waves can be separated by their spindle coupling provides a mechanistic framework to further develop these methods. In particular, we observe that early-fast and late-fast spindles have frequency overlap near the peak frequency for all spindle events, around 14 Hz, and therefore implementation of bandpass filters that share this frequency window may inadvertently mix both early-fast and late-fast spindle events. Given the distinct timing aspects of early-fast and late-fast spindles with regard to their respective slow wave downstate troughs and upstate peaks, this mixing of spindle events may complicate the interpretation of spindle timing measurements.

Aging is associated with decreased amplitude of slow waves,^46^ and changes in slow wave morphology have also been described with increasing age.^55^ The decrement in slow wave amplitude also produces a well-described loss of stage 2 sleep scoring due to technical limitations with use of an absolute slow wave size threshold rule for scoring each segment of NREM sleep as stage N2 vs N3 sleep.^40,46^ These considerations certainly influence the interpretation of our observations demonstrating a shift in coupling subtypes in stage N2 sleep with increasing age. The stage N2 mix of slow wave subtypes is expected to include more N3 slow wave subtypes as age increases due to these sleep stage scoring rules.

In addition to early-fast and late-fast spindles, events occurring prior to the trough of the slow wave downstate include the originally described slow spindles^27,49,50^ and the more recently described theta bursts.^29-31^ While little is known about the function of slow spindles, theta bursts are proposed to trigger slow wave events,^29-31^ and may also precede hippocampal ripples during memory consolidation^29,30^. Challenges of surface EEG complicate the interpretation of theta burst power within our analysis, as baseline segments for EEG normalization contribute to the EEG power calculations of time-frequency map ROIs. The baseline of stage N3 sleep is observed to normalize theta frequencies disproportionally to stage N2 baseline, largely due to frequency leakage from the delta power of N3 sleep. Nonetheless, our data demonstrating a separation of slow wave events by spindle coupling (Figure 6) eliminate this confound by use of a common baseline, and these data do suggest that theta bursts and slow spindles might be more frequently associated with early-fast spindle-coupled slow wave events. Notably, theta bursts typically precede slow waves and K-complexes of stage N2, and are less likely to precede slow waves of stage N3 sleep.^31^ Thus, although somewhat speculative, these observations suggest another possible link to a global (light sleep) vs local (deep sleep) relationship with distinct subtypes of slow wave and spindle coupling.

One of the strengths of our study is the utilization of a large sample size to find consistencies in slow waves and spindle coupling amongst the great individual variability that we and others have observed.^48^ This variability is also a weakness of our approach, in that we remain unable to precisely quantify different subtypes of slow waves and spindles at the single-subject level given variability in both frequency and timing of these events between individuals. Further, our experience aligns with other studies in that we find many spindles and slow waves do not demonstrate coupling when analyzed from surface EEG.^27,44,48^ These limitations remain one of the major barriers to the development of this technique as functional biomarker that could be used to predict neurodegenerative disease within an individual aging adult. Future work will be needed to identify additional defining attributes of slow waves and spindles and to refine individualized sorting methods for more consistent quantification of coupling subtypes.

Another technical aspect to consider is our selection of baseline segments for normalization of EEG power, which is necessary to compare coupling across individuals. Previous studies of coupling have applied different approaches in their method of EEG baseline selection, each with potential strengths and weaknesses.^20,27,48,56^ Here we chose to utilize regions of baseline activity that match the stage of sleep for each respective slow wave population, similar to work by Muehlroth et al,^20^ to allow sleep stage-specific quantitative measurements. Uncoupled spindles are abundant during NREM sleep, and these spindles within baseline segments will impact the measured coupled spindle power when their respective frequencies overlap. This method of baseline selection provides an advantage within our study as it allows clearer delineation of overlapping early-fast and late-fast spindles; however, there is a limitation in the ability to quantify regions within the frequency band common to both coupled and uncoupled spindles. Development of more complex signal processing techniques may provide a normalization method without these potential frequency biases for future studies.

In conclusion, we find that human NREM sleep contains an amalgam of distinct slow wave and spindle coupling events that mix together to form composites of slow wave sleep. Stages N2 and N3 sleep contain differing amounts of early-fast spindles versus late-fast spindles, possibly reflecting distinctions in the function of slow wave subtypes between the stages of sleep. The composition of slow wave and spindle subtypes can be observed to transition across phases of the human life cycle, and a predominant pattern of deeper sleep coupling emerges during the aging process. These insights provide a conceptual foundation to advance our knowledge of sleep’s memory functions. Finally, several clinically relevant advancements may come from these observations in both efforts to utilize slow wave and spindle physiology as a biomarker for detection of early Alzheimer’s disease^22,57,58^ and ongoing clinical trials aiming to restore or enhance slow wave sleep as a clinical intervention.^25,59,60^

## Acknowledgements

The authors are grateful for the open access to data provided by the Cleveland Family Study (CFS) (supported by grants from the National Institutes of Health (HL46380, M01 RR00080-39, T32-HL07567, RO1-46380)), and the National Sleep Research Resource (supported by the National Heart, Lung, and Blood Institute (R24 HL114473, RFP 75N92019R002)).

---

**Supplementary Table 1.** Demographics of the study population subset used to identify regions of interest.

| *Characteristic* | *N* | *N (%) or Median (IQR)* |
| --- | --- | --- |
| Age | 88 | 43.31 (24.43-52.70) |
| Sex | 88 |  |
| - Female |  | 45 (51.14) |
| - Male |  | 43 (48.86) |
| Race | 88 |  |
| - White |  | 35 (39.77) |
| - Black |  | 52 (59.09) |
| - Other |  | 1 (1.14) |
| Apnea Hypopnea Index | 88 | 7.28 (1.59-22.54) |
| Apnea Hypopnea | 88 |  |
| - Less than 15 |  | 57 (64.77) |
| - 15 or greater |  | 31 (35.23) |
| % Time in Stage 1 | 81 | 15.00 (10.00-22.00) |
| % Time in Stage 2 | 80 | 223.00 (184.00-249.00) |
| % Time in Stage 3 | 80 | 16.85 (10.50-25.50) |
| % Time in REM | 83 | 70.00 (50.00-89.00) |

**Supplementary Figure 1:**
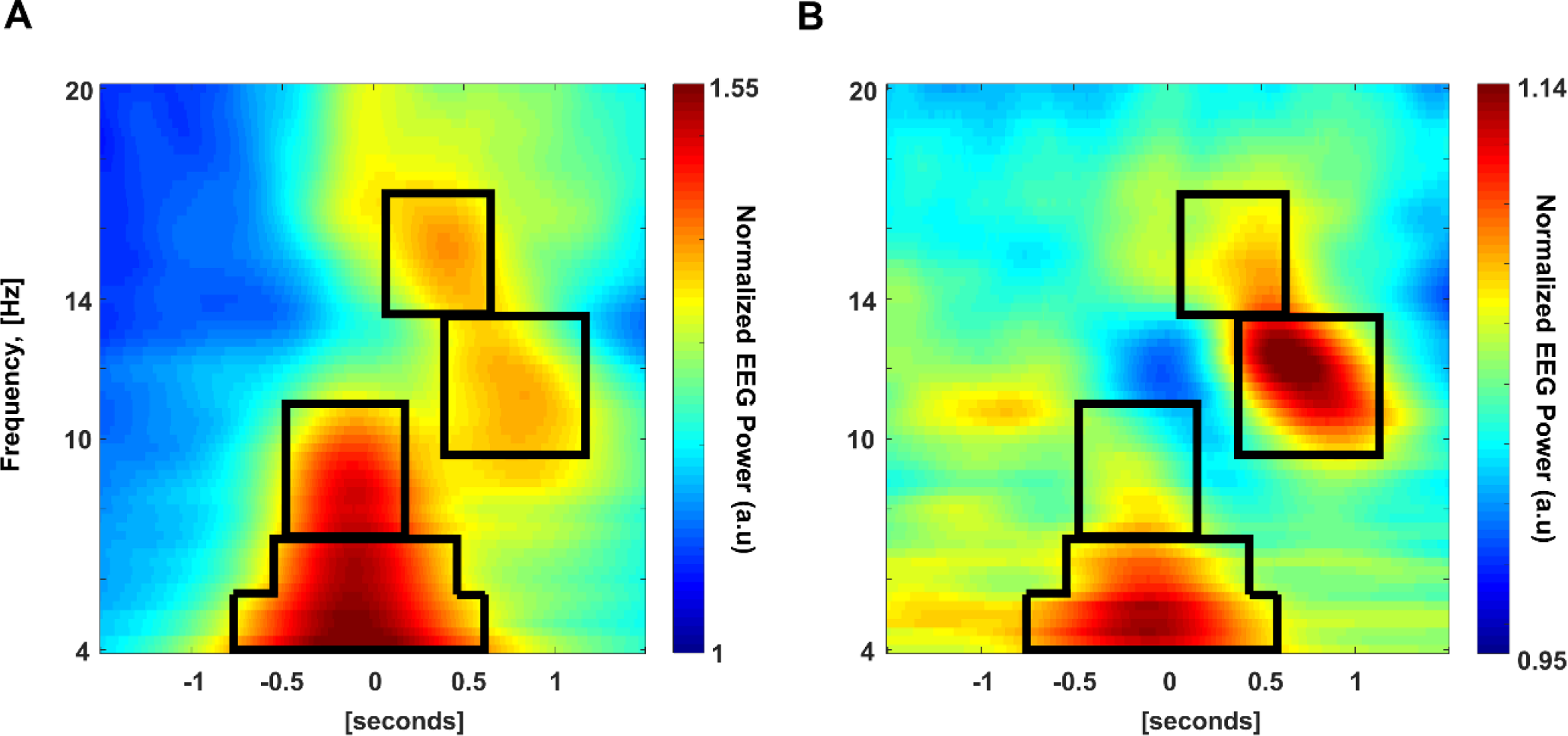
Region of interest selection. Four regions were defined by visual inspection of averaged wavelet plots for 88 subjects in stage N2 sleep (A) and stage N3 sleep (B).

**Supplementary Figure 2:**
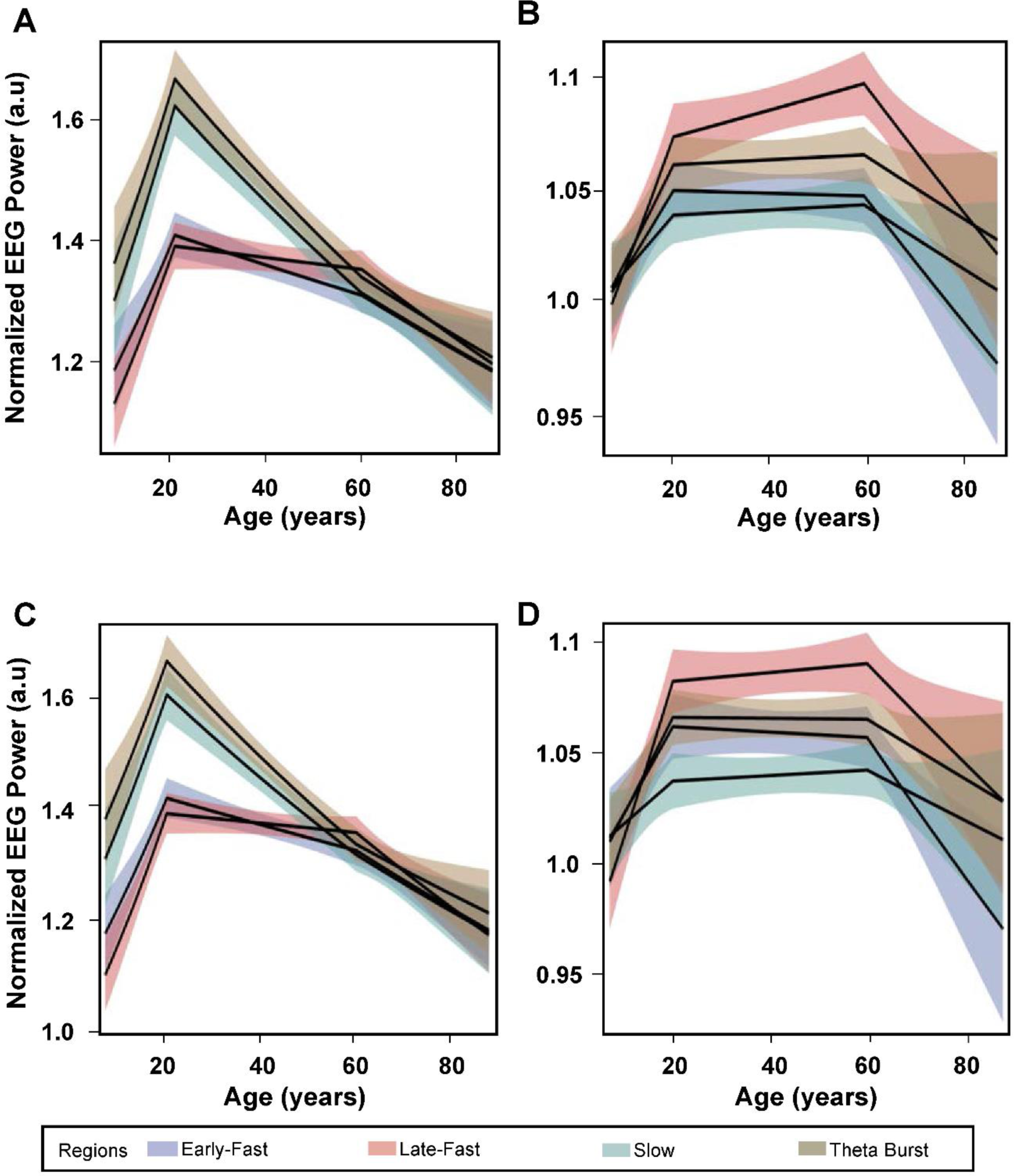
Mixed model estimates of normalized EEG power (arbitrary units) for each ROI across all subjects with 95% confidence intervals in color shading. Electrode C3 values for stage N2 sleep (A) and stage N3 sleep (B). Electrode C4 values for stage N2 sleep (C) and stage N3 sleep (D).

